# Capturing sclera anisotropy using direct collagen fiber models. Linking microstructure to macroscopic mechanical properties

**DOI:** 10.1101/2024.09.12.612702

**Authors:** Fengting Ji, Mohammad R. Islam, Frederick Sebastian, Xuehuan He, Hannah Schilpp, Bingrui Wang, Yi Hua, Rouzbeh Amini, Ian A. Sigal

## Abstract

Because of the crucial role of collagen fibers on soft tissue mechanics, there is great interest in techniques to incorporate them in computational models. Recently we introduced a direct fiber modeling approach for sclera based on representing the long-interwoven fibers. Our method differs from the conventional continuum approach to modeling sclera that homogenizes the fibers and describes them as statistical distributions for each element. At large scale our method captured gross collagen fiber bundle architecture from histology and experimental intraocular pressure-induced deformations. At small scale, a direct fiber model of a sclera sample reproduced equi-biaxial experimental behavior from the literature. In this study our goal was a much more challenging task for the direct fiber modeling: to capture specimen-specific 3D fiber architecture and anisotropic mechanics of four sclera samples tested under equibiaxial and four non-equibiaxial loadings. Samples of sclera from three eyes were isolated and tested in five biaxial loadings following an approach previously reported. Using microstructural architecture from polarized light microscopy we then created specimen-specific direct fiber models. Model fiber orientations agreed well with the histological information (adjusted R2’s>0.89). Through an inverse-fitting process we determined model characteristics, including specimen-specific fiber mechanical properties to match equibiaxial loading. Interestingly, the equibiaxial properties also reproduced all the non-equibiaxial behaviors. These results indicate that the direct fiber modeling method naturally accounted for tissue anisotropy within its fiber structure. Direct fiber modeling is therefore a promising approach to understand how macroscopic behavior arises from microstructure.

## 1. Introduction

Collagen fibers serve as the primary load-bearing component of soft tissues, and in particular of sclera [1–5], Hence, they are crucial for understanding the mechanical behavior of the eye, and how it relates to eye physiology and pathology [4, 6–8]. Because of the fragility and difficulty in accessing the eye, especially the posterior pole, computational modeling has emerged as a key approach to study the relationship between scleral fiber structure and mechanics [5, 9–14]. The most popular techniques for modeling sclera, however, are based on a continuum mechanics framework and do not account for several potentially crucial characteristics such as fiber interweaving, fiber-fiber interactions and the long-distance effects of fibers. Neglecting interweaving and fiber-fiber interactions can, for example, lead to significant errors when estimating the mechanical properties of sclera fibers by inverse fitting [15]. Accurately incorporating fiber characteristics at the microscale is also likely crucial to predicting correctly the effects at the cellular and axonal scales [16].

We recently introduced a direct fiber modeling technique [17]. The approach is based on detailed histology-based specimen-specific fiber orientations obtained through polarized light microscopy (PLM) [18] while considering fiber interweaving, fiber-fiber interactions, and long fibers. We have demonstrated the direct fiber modeling approach in two studies with different scales. In the first we reconstructed a model of a small sample of temporal sclera, and then used an inverse fitting approach to match experimental equi-biaxial stress-strain data from the literature, simultaneously capturing the behavior in both radial and circumferential directions [17]. In the second study, we applied the direct fiber modeling approach to reconstruct a model of the optic nerve head and adjacent tissues, a substantially larger region than the sclera sample. Because of the size and complexity of the region modeled we had to modify how we used the histological data to reconstruct the model. For example, we focused on modeling fiber bundles 200-400um-thick over small fibers. The model successfully captured the gross behavior of the optic nerve head under inflation caused, including the emergent nonlinear behavior despite having been simulated with linear tissue mechanical properties [19]. Although both studies show the promise of the direct fiber modeling approach, they have important limitations. First, the histological information was from a sheep eye and the experimental tests data from a pig eye. Second, only one posterior sclera sample was analyzed, and therefore we could not establish that the technique can recover specimen-specific information. Third, our previous testing of the model’s behavior was limited to equi-biaxial loading conditions, neglecting the more complex anisotropic conditions that the sclera may experience in physiological contexts [2].

Our aim in this study was to conduct a much more challenging test of the direct fiber modeling’s approach ability to capture sclera microstructure and anisotropic mechanics. First, we evaluated the accuracy of a direct fiber model representation of the complex specimen-specific collagen structure of four sclera samples from different eyes. Second, we evaluated the ability of a direct fiber model to capture specimen-specific anisotropy under equibiaxial and non-equibiaxial conditions. This work helps understand better the capabilities of direct fiber modeling of sclera so that the tool can later be used to study other soft fibrous tissues.

## 2. Methods

This section is organized in three parts according to the project steps. First, biaxial mechanical testing was conducted on healthy sheep posterior pole samples to obtain their stress-strain responses under five different loading protocols. Second, the same samples used for mechanical testing were fixed and sectioned, and the sections imaged using PLM. The PLM images were post-processed and used to build direct fiber models with the fiber orientation data. Each direct fiber model consisted of a fibrous component embedded in a matrix representing the non-collagenous components. The combined fiber and matrix model was used in an inverse fitting optimization process to match the simulated stress-strain behaviors with experimental data acquired under equi-biaxial testing. The inverse process produced specimen-specific fiber material properties and pre-stretching strains.

All the image processing was done in MATLAB v2020 (MathWorks, Natick, MA, USA) and FIJI is Just ImageJ (FIJI) [20, 21]. Modeling was done in Abaqus 2020X (Dassault Systemes Simulia Corp., Providence, RI, 171 USA). Customized code and the GIBBON toolbox [22] for MATLAB v2020 (MathWorks, Natick, MA, USA) were used for model pre/post-processing and inverse fitting.

### 2.1 Biaxial mechanical testing

The experimental procedures were conducted in Northeastern University, utilizing a methodology previously described [23, 24]. The study was conducted in accordance with the tenets of the Declaration of Helsinki and the Association of Research in Vision and Ophthalmology’s statement for the use of animals in ophthalmic and vision research. Three fresh sheep eyes were obtained from a local slaughterhouse within 24 hours postmortem and immediately transferred to the lab in a cold isotonic phosphate buffer saline (PBS) solution. Upon arrival, two posterior sclera samples measuring 11mm x 11mm were carefully excised from both temporal and superior quadrants, approximately 2 to 3 mm away from the sclera canal (Figure 1 A). For the purpose of optical tracking and tissue deformation/strain analysis, four submillimeter glass markers were affixed to the surface of each sample.

**Figure 1.**
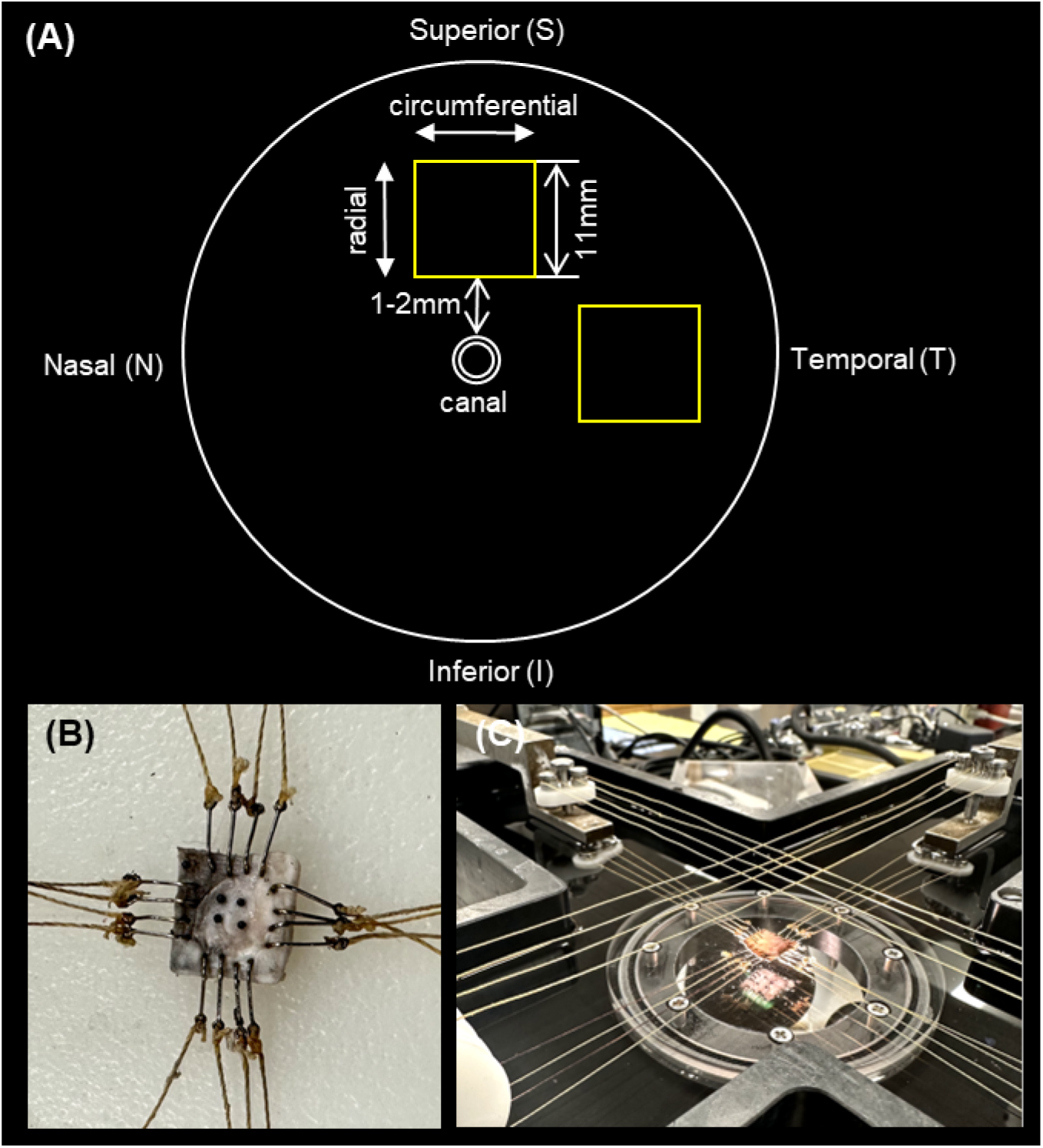
Key aspects of the experimental setup and procedures for the biaxial testing. (A) Schematic diagram showcasing the posterior pole of the eye, highlighting the locations of the two samples (depicted as yellow boxes) that were obtained specifically for the biaxial testing. (B) An example image showcasing a sample with hooks attached, ready for biaxial testing. The hooks ensure proper mounting and fixation of the sample during the experimental procedures. (C) The sample was mounted on a custom-built biaxial mechanical testing system. The loading axes of the system were aligned with the circumferential and radial directions of the sample, ensuring precise application of stress in the desired directions. The sample was then subjected to the five loading protocols. Note that the crossed strings, as depicted at the lower right of the sample, were corrected prior to conducting the experiment.

The sample was then mounted onto the biaxial testing equipment using fish hooks, with the loading axes aligned with the circumferential and radial directions of the sample (Figure 1 B and C). A 0.5g tare load was initially applied to flatten the sample. Subsequently, each sample was subjected to biaxial stress control and underwent five distinct loading protocols (Table 1). Each loading protocol consisted of ten 20-second cycles. The first nine cycles served as a preconditioning phase, while data from the last loading cycle was used for subsequent analysis. Based on preliminary tests, a maximum stress level of 150kPa was applied, as it allowed the sample to maintain sample shape without incurring damage. Once the stress-strain data were obtained, the samples were carefully unmounted from the biaxial testing equipment and immersion fixed in 10% formalin for 24 hours. We chose formalin because it has been shown to cause only minimal changes in the shape or size of ocular tissues [25].

**Table 1.** Maximum radial (anterior-posterior) and circumferential (equatorial) stress values for each biaxial loading protocol.

| Loading Protocol # | Radial (kPa) | Circumferential (kPa) |
| --- | --- | --- |
| 1:1 | 150 | 150 |
| 1:0.75 | 150 | 112.5 |
| 0.75:1 | 112.5 | 150 |
| 1:0.5 | 150 | 75 |
| 0.5:1 | 75 | 150 |

Temporal and superior samples from three eyes produced six samples. Two of the samples were excluded from further analysis because of tissue damage incurred during the experimental procedures. Hence, the rest of the analysis is based on stress-strain responses from four samples. More details regarding these samples can be found in Table 2.

**Table 2.** Information about the four posterior sclera samples used for direct fiber modeling. The samples were obtained from both the superior and temporal quadrants of three sheep eyes. Each sample had a different thickness, resulting in a variation in the number of collected coronal sections.

| Sample # | Temporal/Superior | Sheep # | Thickness($\mu\text{m}$) | Number of Coronal Sections |
| --- | --- | --- | --- | --- |
| 1 | Superior | Eye-1 | 528 | 50 |
| 2 | Temporal | Eye-2 | 1168 | 87 |
| 3 | Superior | Eye-2 | 635 | 60 |
| 4 | Temporal | Eye-3 | 838 | 74 |

At the initial stages of the loading tests, the experimental stress-strain data exhibited relatively high noise and variability compared with more stable later steps. These points were excluded from the inverse fitting process. We applied a moving average smoothing algorithm to reduce the noise in the rest of the stress-strain curves [26]. For the rest of the analysis, we used the experimental data after smoothing.

### 2.2 Histology, polarized light microscopy and image post-processing

Following the fixation process, the samples were cryo-sectioned into 20 μm slices (Figure 2). To be able to reconstruct accurately the 3D architecture of the samples we used a process involving both coronal sections (parallel to the surface of the sample) and sagittal sections (perpendicular to the surface sample). For coronal sections, serial sectioning was performed at the center of the sample, every section was collected starting when there was visible sclera and stopping when the sclera was no longer visible. The number of sections collected varied among the different samples, and the specific quantities are detailed in Table 2. For sagittal sections, two slabs from the edge of the square-shaped sample were obtained. In this way we were able to obtain high resolution coronal data from the core of the tested sample, and high-resolution transverse data at the core edge. Coronal and sagittal sections approximate two orthogonal views of the collagen structure, providing information on the three-dimensional organization of the fibers.

**Figure 2.**
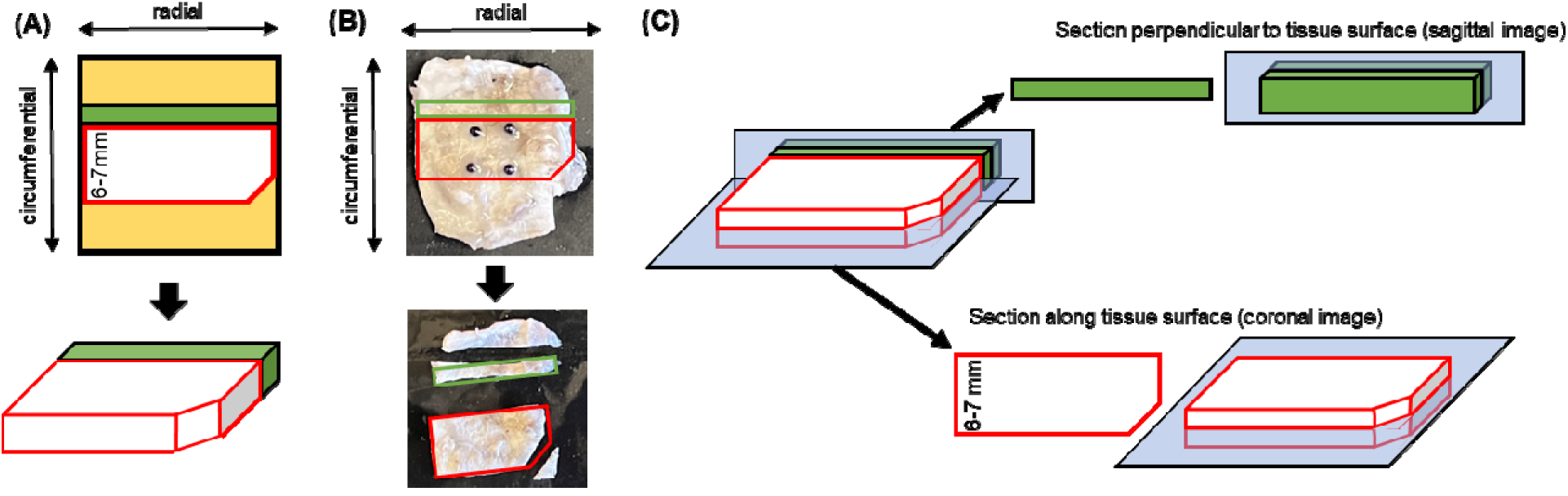
Process of sectioning a sclera sample. (A) Sclera sample after biaxial testing was processed for sectioning (top: 2D view; bottom: 3D view of the region for sectioning). A notch at the corner of the sample was used to indicate the tissue directions. Since it was not feasible to section the same piece of tissue both coronally and sagittally, a sample (depicted as a white block, the shorter length was 6-7mm) was obtained from the center of the tissue. Additionally, another sample was acquired from the adjacent tissue next to the white block (shown as a green block). (B) An example image of the sclera tissue with the two samples dissected. Prior to dissection, the fiberglass markers attached to the tissue were carefully removed. (C, top) The green block was sectioned sagittally, resulting in sagittal sections of the tissue. (C, bottom) The white block was coronally sectioned, allowing for the acquisition of serial sections without any loss. The blue surface depicted in the image represented the plane of sectioning.

All sections were imaged with PLM as described before [18, 27]. Briefly, two polarized filters (Hoya, Tokyo, Japan) were used, one a polarizer and the other an analyzer, to collect images at four filter orientations 45° apart. The images were all captured using an Olympus MVX10 microscope (1× magnification setting, 6.84 µm/pixel).

PLM images were processed to derive the in-plane collagen orientation at each pixel (in Cartesian coordinates) and a parameter that we referred to as “energy” [28]. Energy helped identify regions without collagen, such as outside of the section, and regions where the collagen fibers were primarily aligned out of the section plane, so that they can be accounted for in the orientation distribution. Following the processing of PLM images, all images obtained from a single sample were sequentially stacked and registered based on the sharp edges [29]. After registration, the original images underwent reprocessing to obtain “corrected” orientation angles that are consistent across all the sections.

To focus on a specific area for subsequent construction of the direct fiber model, we selected a square-shaped block positioned at the center of the coronal sections. The dimensions of this selected block were 4.1x4.1mm (Figure 3A). However, due to the irregular shape of the tissue and the folds caused during tissue sectioning, the stack of cropped blocks exhibited variations in thickness (Figure 3C). In order to facilitate fiber tracing for the construction of the direct fiber model, we implemented a linear interpolation algorithm, as described by Akima [30, 31]. The interpolation algorithm was employed to interpolate angle and energy values at each location in-depth within the stack, thereby converting the inconsistent thicknesses into a uniform value. As a result of this interpolation, the image stack was transformed into a uniform and regular block (Figure 3D). Importantly, this interpolation process did not alter the orientation distribution of the fibers within the stack.

**Figure 3.**
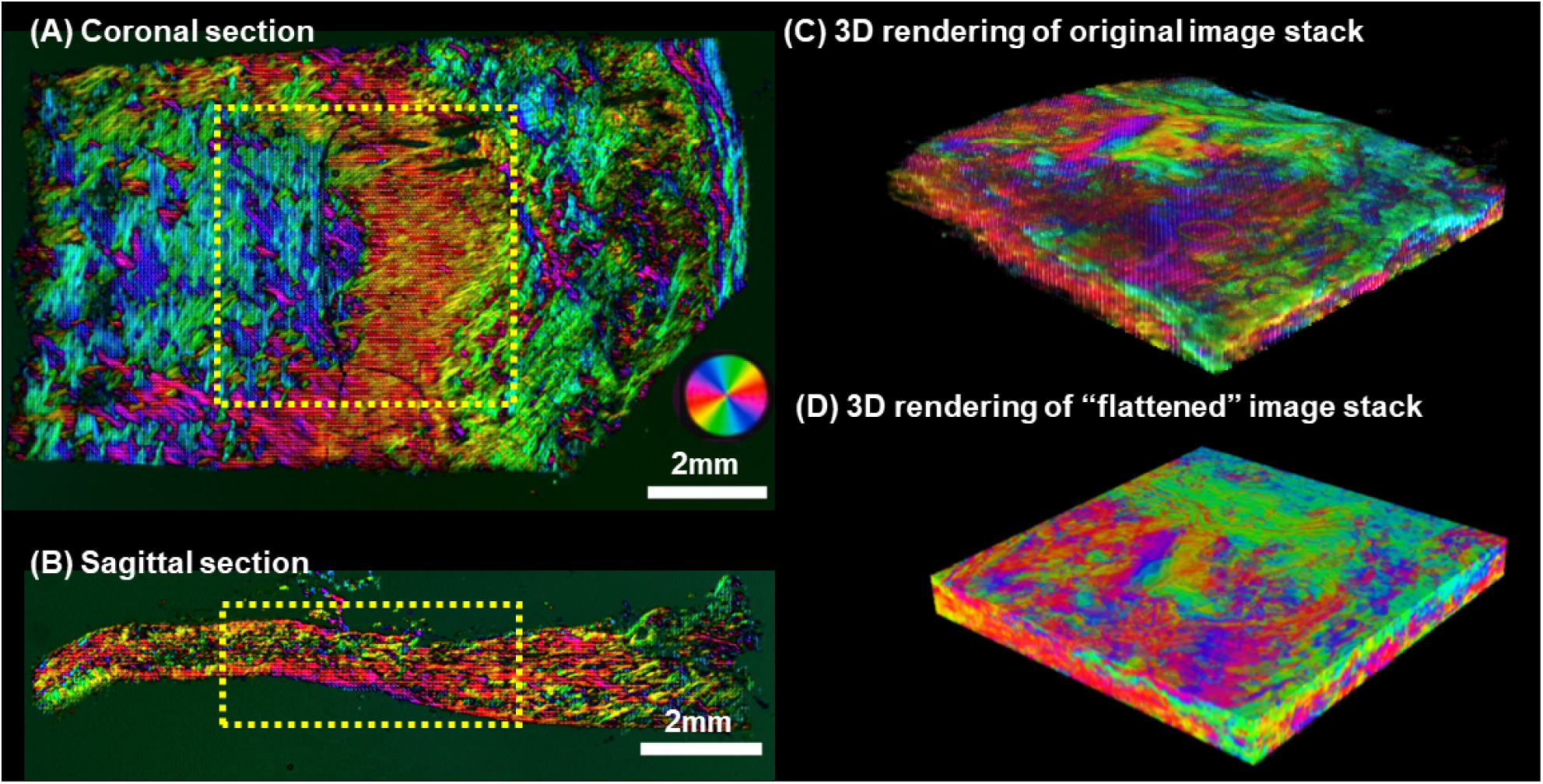
Example PLM images and process of image post-processing. (A) Example PLM image of a coronal section. A square-shape block was cropped from the stack and used as a reference to build the direct fiber model. (B) Example PLM image of a sagittal section. The model’s in-depth fiber orientation distribution was adjusted based on the orientations obtained from the yellow block. The position of this block corresponded to the position of the square-shaped block obtained from the coronal section. The colors indicate the local fiber orientation in the section plane, with brightness proportional to the “energy” parameter. (C) The original stack showed irregular thickness from one location to another. (D) After interpolation, the thickness of the stack was uniformized which facilitates the tracing of fibers during the construction of the direct fiber model.

To validate the geometry of the direct fiber model, we computed both the coronal and sagittal collagen fiber orientation distributions. For the coronal orientation distribution, we utilized the pixel-level PLM data obtained from the original image stack, taking into account the local “energy” at each pixel. Similarly, for the sagittal orientation distribution, we summed up the PLM data obtained from the cropped block of the two sagittal sections (depicted in Figure 3B). Again, the energy weighting was applied to ensure accurate representation of the collagen fiber orientations in the sagittal plane.

### 2.3 Direct fiber modeling

#### 2.3.1 Model construction

We constructed four models (Figure 4A) based on the procedure described previously [17], corresponding to each sample listed in Table 2. Briefly, fibers were simulated using 3- dimensional linear truss elements (T3D2 in Abaqus). The locations of fibers were defined by sampling orientation values from PLM images at regularly spaced “seed” points (437 μm apart).

**Figure 4.**
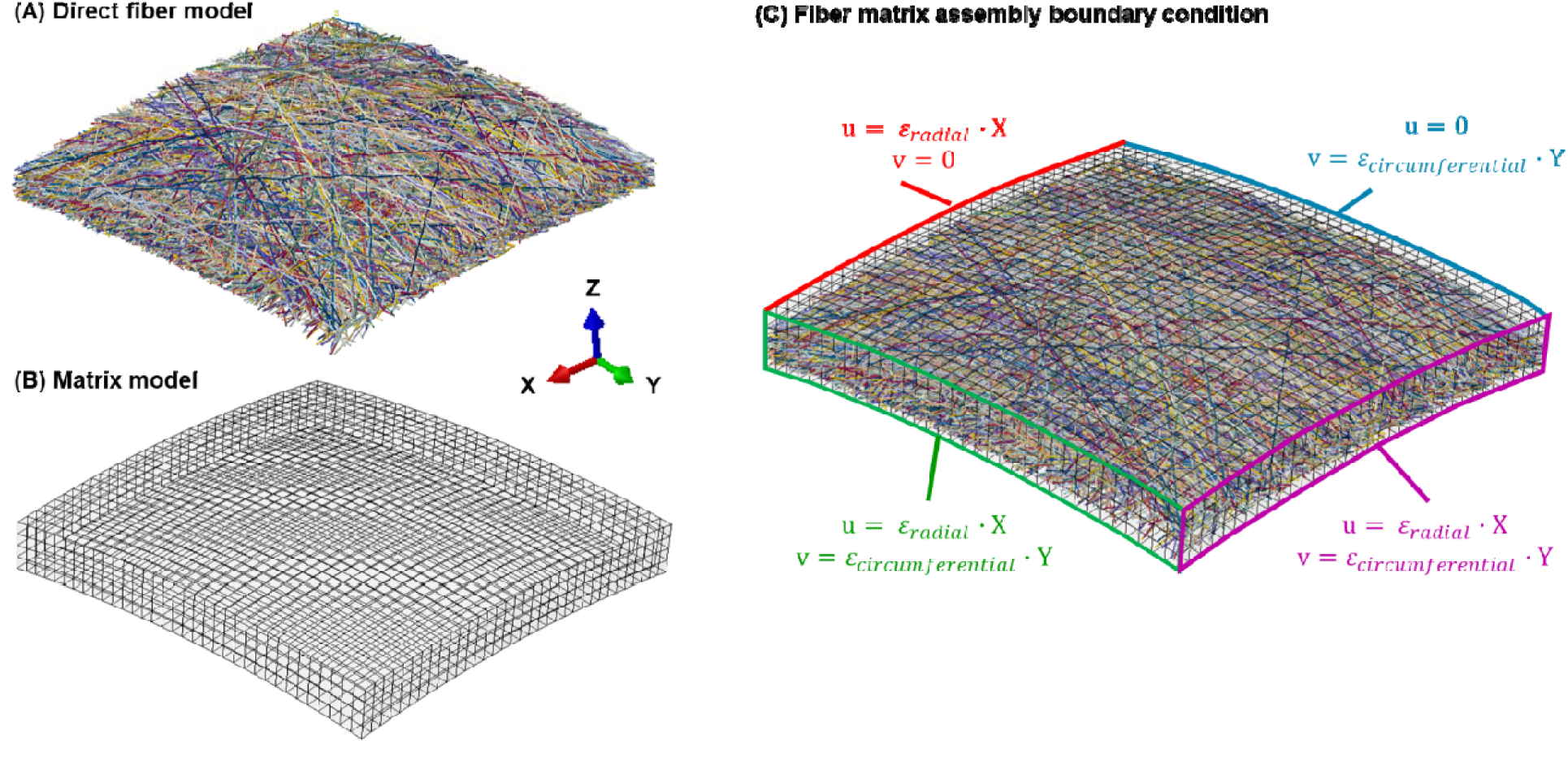
Isometric view of an example (A) direct fiber model and (B) matrix model. (C) Displacement boundary conditions were applied to the fiber matrix assembly, with components u, v representing displacement in X (radial) and Y (circumferential) direction, respectively. The displacement in Z direction was not constrained, given that fish hooks and strings did not restrict the sample’s displacement in the Z direction. In the first step biaxial stretch, the value of ε_radial_ and ε_circumferential_ were optimized and derived in the inverse fitting procedure. In the second step biaxial stretch, the experimental strain values were assigned to εradial and εcircumferential, aiming to match the model with the stress-strain behaviors observed in the experimental data.

Straight fibers, 13.68 μm in diameter, were traced at each seed point based on its orientation angle. The process was repeated for each layer, resulting in a stack of 2D layers with interpenetrating fibers. An algorithm was employed to resolve interpenetrations by iteratively shifting the elements until they no longer overlapped [32, 33]. Fiber elements were re-meshed to maintain lengths between 82μm and 164μm, while controlling the minimum radius of curvature for smoothness. The amplitudes of the fiber undulations in-depth were adjusted to more accurately represent the distribution of fibers in three dimensions.

To account for the natural curvature of the sclera, an additional step was performed to adjust the flat fiber model and match it to the curvature of the eyeball. The external radius of three sheep globes was manually measured, and the average radius was determined to be 14856 μm. The flat model was then projected onto a sphere with an external radius of 14856 μm, effectively introducing curvature to the model. It was important to note that this implied assuming that the sheep eye locally resembled a sphere.[34] To assess the similarity between the model and the PLM images, the orientation distribution of the curved model was quantified and compared with the distribution obtained from the PLM images. This comparison aimed to evaluate how well the model captured the observed fiber orientations in the images. The quantification involved counting the occurrences of element orientations within the model, where each element’s orientation represented the slope angle in the section plane. This approach accounted for varying element sizes and enabled a proper comparison with the pixel-based measurements obtained from PLM. To evaluate the fitness of the orientation distributions, adjusted R-squared (adjusted R^2^) values were employed [35]. The adjusted R^2^ values provided a measure of how well the model’s orientation distribution fit the distribution observed in the PLM images. The use of adjusted R^2^ values allowed for the consideration of the complexity of the model and the number of parameters involved, providing a more robust evaluation metric than a conventional R^2^.

In addition to the fiber model, a matrix model was also constructed (Figure 4B). The matrix model was designed to have the same dimensions and shape, including the same curvature, as the fiber model, ensuring consistency between the two. The end-nodes of the fibers were positioned on the surfaces of the matrix, creating a cohesive representation of the fiber-matrix structure.

#### 2.3.2 Model inverse fitting

##### 2.3.2.1 Meshing and material properties

Fibers were modeled as a hyperelastic Mooney-Rivlin material [36]:

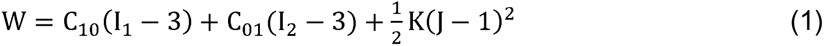

where W was the strain energy density, C_1O_ and C_O1_ were the material constants, restricted by C_1O_ + C_O1_ > 0 and would be determined by inverse fitting, I_1_ and I_2_ were the first and second invariants of the right Cauchy-Green deformation tension, K was the bulk modulus and J was the determinant of the deformation gradient. The matrix was modeled as a neo-Hookean material with a shear modulus of 200 kPa [3, 37].

The fiber components were modeled using 3-dimensional linear truss elements (T3D2) in Abaqus. The length of the fiber elements ranged from 82 μm to 164 μm, resulting in aspect ratios between 6 and 12. The matrix was meshed using linear eight-noded hybrid hexahedral elements (C3D8H) in Abaqus. The element size for the matrix varied among the samples due to differences in sample thickness, with 4 to 6 elements spanning the shell thickness. To ensure model accuracy, a mesh refinement study was conducted. The fiber model’s mesh density was doubled, while the matrix model had its mesh density doubled in both in-plane directions and the thickness direction. The study’s findings indicated that altering the mesh density had a negligible impact on stress predictions, with maximum stress values changing by less than 1%. Based on these findings, the chosen mesh density was deemed sufficient to ensure numerical accuracy in the obtained results.

##### 2.3.2.2 Interactions

Fiber-fiber interactions were simulated in the following two ways. First, the interactions were considered by preventing fiber interpenetrations using Abaqus’ general contact with no friction. Second, to enhance the interweaving effect of fibers and prevent them from sliding apart during stretching, a method involving the constraint of nodes was employed. Approximately 10% of the nodes, primarily located in the outer surface or boundary of the model, were selected. These nodes were connected to their closest neighboring nodes, and their relative motion in the Z direction was constrained to zero using linear constraint equations in Abaqus. By applying these constraints, the free ends of the fibers were better controlled, resulting in a more stable and realistic model.

Fiber-matrix interactions were ignored, as is usual in biomechanical models of the eyes [5, 38, 39].

##### 2.3.2.3 Finite element analysis procedure

The fiber-matrix assembly underwent a quasi-static biaxial stretching process to match the experimental stress-strain data obtained in Method Section 2.1. The matrix was simulated using Abaqus standard implicit procedure, while the direct fiber model utilized Abaqus dynamic explicit procedure to enhance convergence and computational efficiency. The resulting stresses σ in the radial and circumferential directions were a combination of matrix and fiber contributions, with the matrix stress weighted by the fiber volume fraction (VF) as shown in Equation (2):

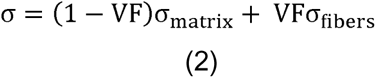

To ensure efficient dynamic analysis, mass scaling was implemented, allowing for a stable time increment of 1e-5. The simulation was conducted within a time period where inertial forces remained negligible, with the maximum kinetic energy kept below 5% of the internal energy to confirm the insignificance of inertial effects.

##### 2.3.2.4 Boundary conditions and inverse fitting procedure

In the biaxial stretching process of the fiber-matrix assembly (Figure 4C), a two-step approach was employed to simulate the experimental conditions. In the first step, biaxial stretching was applied to pre-stretch the model. This step aimed to simulate the initial stretching that occurs when a 0.5g tare load was applied to flatten the tissue. The resulting stress and strain values from this step were not recorded, as they were not included in the reported experimental stress-strain data. In the second step, the model was further stretched, and the same strains observed in the experiment were assigned as the displacement boundary condition to the fiber-matrix model. In this step, the model stress and strain values were recorded, starting from 0. This approach was consistent with the experimental setup, where the stress and strain resulting from the tare load were not included in the reported experimental stress-strain data. By applying this two-step biaxial stretch with the appropriate displacement boundary conditions, the model aimed to replicate the mechanical behavior observed in the experiment and allow for a comparison between the model’s stress-strain behaviors and the experimental data.

During the inverse modeling procedure, there were four parameters to determine. The first two parameters were the pre-stretching strains along the radial and circumferential directions in the first step of the biaxial stretch. Since the resultant strains caused by the tare load were not characterized in the experiment, these values were unknown and needed to be determined through the inverse modeling process. The second two parameters were the material properties of the fibers (C_1O_ and C_O1_).

In the inverse fitting procedure, two optimizations were performed to determine the optimal parameters and match the stress-strain data of the five loading protocols. In the first optimization, all four parameters (pre-stretching strains and fiber material properties) were optimized to match the stress-strain data obtained using loading protocol 1:1. Further discussion on the selection of loading protocol 1:1 can be found in the Discussion section. A simplex search method was used for this optimization [40]. The algorithm aimed to find the values of the parameters that minimized the residual sum of squares (RSS) between the simulated and experimental stress-strain curves. The optimization process continued until the adjusted R^2^ value between the curves exceeded 0.9, indicating a good fit between the model and experimental data. This optimization allowed for the determination of the optimal pre-stretching strains and fiber material properties (C_1O_ and C_O1_).

In the second optimization, only the two pre-stretching strains were optimized, using the derived fiber material properties from the first optimization. This optimization aimed to match the stress-strain data obtained from the remaining four loading protocols. The same search method as before was employed, and the optimization was concluded when the adjusted R^2^ value between the stress-strain curves exceeded 0.9 in both loading directions.

By performing these two optimizations, the model was able to find the optimal parameters that resulted in stress-strain curves closely aligned with the experimental data for all the five loading protocols.

### Preliminary analysis

During the preliminary analysis of the inverse fitting procedure using Sample #1, it was observed that the experimental data from loading protocol 0.5:1 exhibited negative strains along one loading direction. This suggested the presence of anisotropic mechanical properties, with one loading direction being much softer than the other direction. When assigning negative strains to the direct fiber model, it led to instability in the model. This was problematic because the direct fiber model is capable of accurately representing fiber tension, but not as well longitudinal fiber compression and buckling. As a result, all the loading protocols that showed negative strains were excluded from the analysis. Based on the mechanical testing results, the loading protocol 0.5:1 of all the samples presented negative strains (Supplemental data - Figure 1). Therefore, the results related to loading protocol 0.5:1 were not reported in this study. The focus was placed on the remaining loading protocols that did not exhibit negative strains.

## 3. Results

Figure 5 presents the fiber orientation distributions of the direct fiber model and corresponding PLM images for the four samples. The agreement between the model and PLM images was observed in both the coronal and sagittal planes. The alignment of these orientation curves yielded adjusted R^2^ values exceeding 0.89 in all cases, indicating a strong match between the model and experimental data.

**Figure 5.**
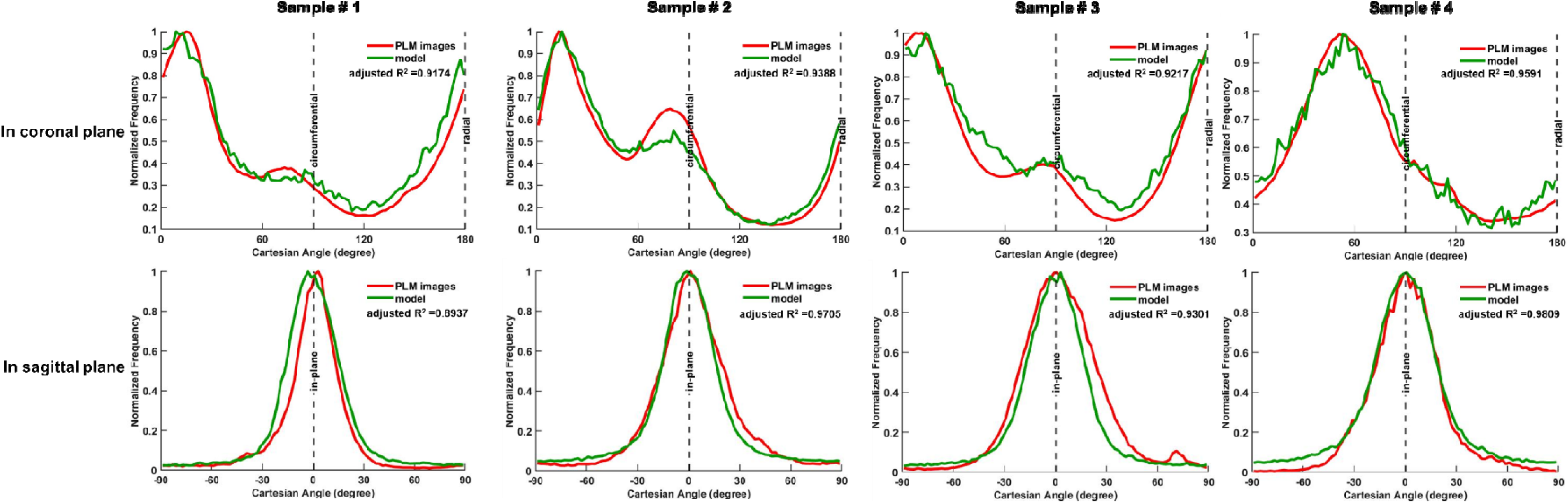
Fiber orientation distributions of the direct fiber model (green lines) and the corresponding PLM images (red lines). The analysis was performed in both the **(top row)** coronal and **(bottom row)** sagittal planes of the four samples. In the coronal plane, the PLM orientation was determined by analyzing a stack of all coronal images, with the radial direction represented by 0 and 180°, and the circumferential direction represented by 90°. The sagittal plane displayed the average orientation distribution obtained from two sections, with the in-plane direction represented by 0°. The frequencies of fiber orientations have been normalized by the total sum of frequencies for effective comparison. The results demonstrate a strong agreement between the fiber orientation distributions of the direct fiber model and those observed in the PLM images, in both the coronal and sagittal planes. All the adjusted R^2^ values, which exceeded 0.89, indicate a high level of similarity between the model and experimental data.

Figure 6 shows fiber displacements and stresses at full stretch of an example sample (Sample #4). The visualization revealed heterogeneous behaviors of the fibers, with varying deformations and stress distributions at the microscale. The model effectively captured the non-uniform response of the fibers under applied loading conditions, highlighting the intricate mechanical behavior within the tissue.

**Figure 6.**
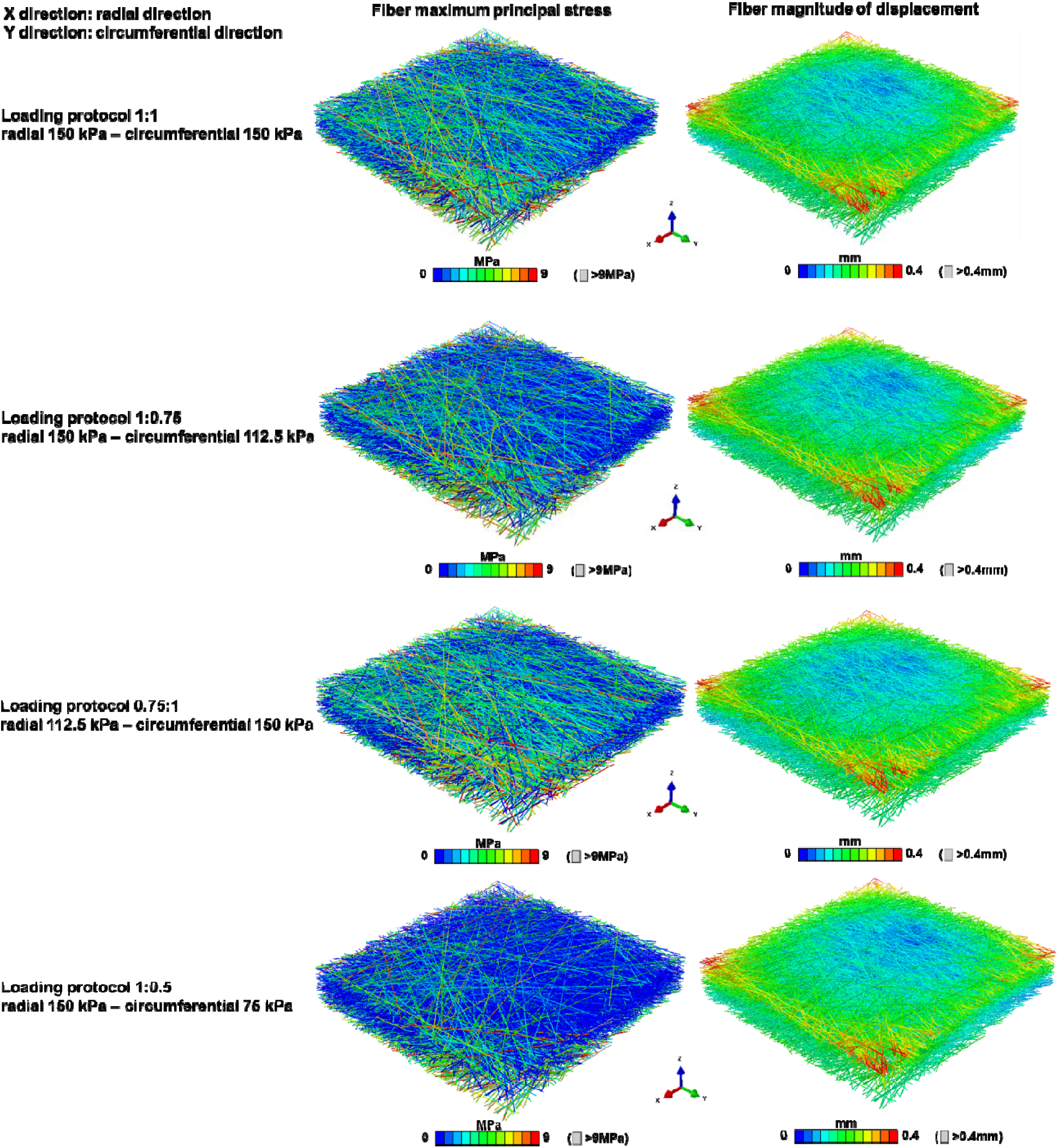
Isometric views of the direct fiber model of Sample #4 at full stretch. The visualization was enhanced by coloring the fibers based on two important mechanical parameters: the maximum principal stress (shown in the left column) and the magnitudes of displacement (shown in the right column) for each loading protocol. The variations in stress and displacement patterns can be observed, highlighting the non-uniform distribution of stresses and displacements within the tissue.

Isometric views of the Sample #4 direct fiber model during loading protocol 1:1 are shown in Figure 7. The visualization uses color to represent the maximum principal stress (left column) and displacement magnitudes (right column). Initially, the model showed curvature, which gradually flattened during stretching. As expected, more fibers experienced higher stress as stretching was applied, in a process of recruitment.

**Figure 7.**
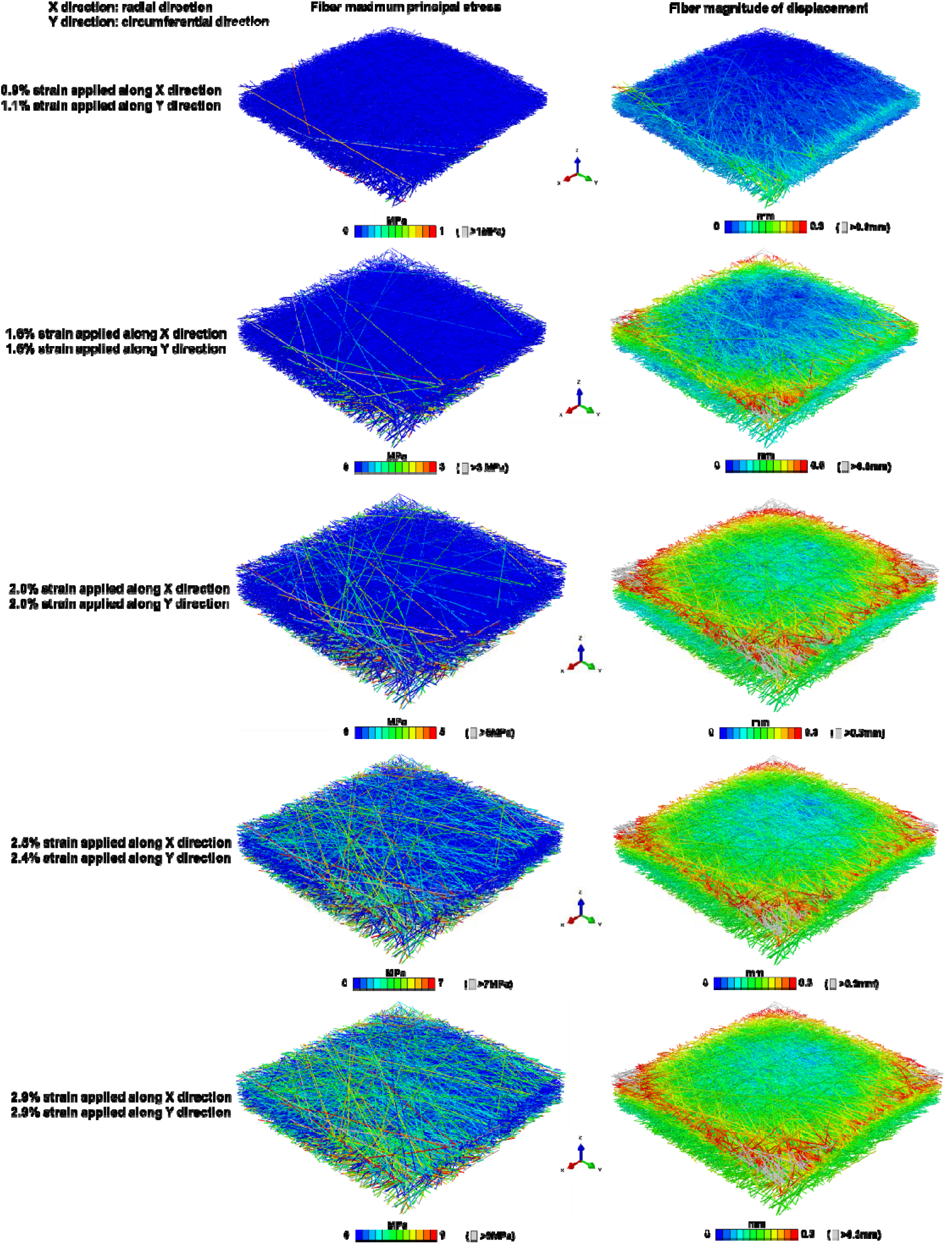
Isometric views of the direct fiber model of Sample #4 at different stages of the simulation while undergoing loading protocol 1:1. The visualization was colored based on the maximum principal stress (left column) and the magnitudes of displacement (right column). In the early stage of the simulation, the model exhibited some curvature. As it underwent stretching, the model gradually transformed into a flattened configuration. As stretching was applied, a larger number of fibers experience higher levels of stress.

Stress-strain curves of the optimized models are shown in Figure 8, showcasing the excellent agreement with experimental data in both the radial and circumferential directions for most of the loading protocol cases. The majority of adjusted R^2^ values exceeded 0.9, indicating a robust fit between the model predictions and experimental observations. The derived parameters that achieved this high level of agreement were provided in Table 3.

**Figure 8.**
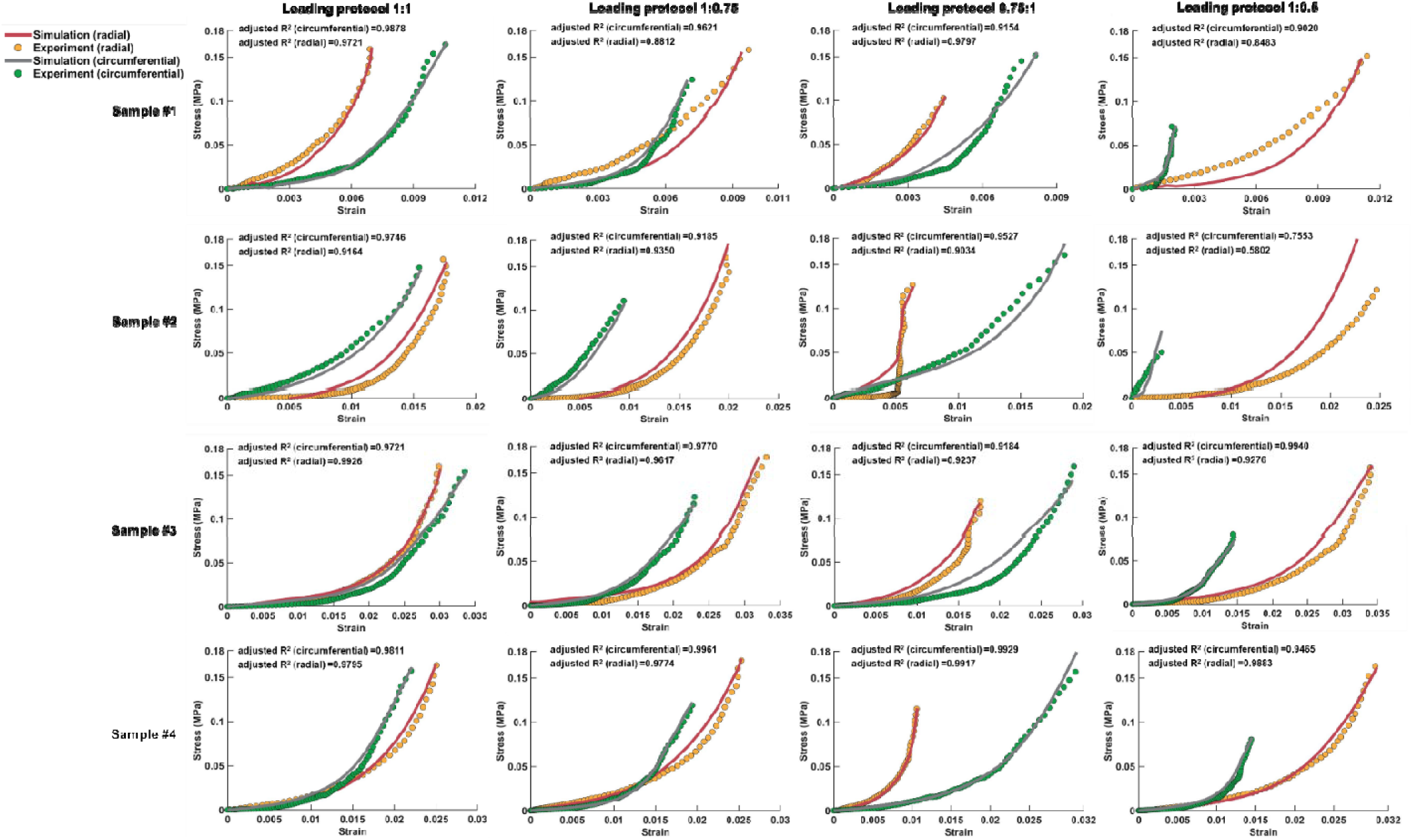
Stress-strain responses of the fiber-matrix assembly and the corresponding experimental data. The stress-strain curves were presented for both the radial and circumferential directions in each of the loading protocols. The fiber material properties were determined by fitting the model to the stress-strain data of loading protocol 1:1 (first column). These derived material properties were subsequently utilized directly for the inverse fitting in other loading protocols. The results demonstrated a successful fit between the model and the experimental data, with consistent agreement achieved simultaneously in both the radial and circumferential directions for each loading protocol. The goodness of fit was quantified using the adjusted R^2^ value, which exceeded 0.9 in most of the cases. Notably for each sample, the stress-strain responses were obtained using the same fiber material properties throughout the simulations, with the variations observed solely in the pre-stretching strains.

**Table 3.** The optimized fiber material properties (C_10_ and C_01_) and the pre-stretching strains along radial and circumferential directions.

| Sample # | Loading protocol # | $C_{10}$ | $C_{01}$ | Pre-stretching strain<br>(radial) | Pre-stretching strain<br>(circumferential) | Fiber elastic modulus<br>(GPa) |
| --- | --- | --- | --- | --- | --- | --- |
| 1 | 1:1 | 4030 | -2.0150 | 0.00825 | 0.00475 | 23.97 |
|  | 1:0.75 |  |  | 0.00600 | 0.00725 |  |
|  | 0.75:1 |  |  | 0.00950 | 0.00750 |  |
|  | 1:0.5 |  |  | 0.00450 | 0.01050 |  |
| 2 | 1:1 | 542.7 | -0.1740 | 0.00050 | 0.00775 | 3.22 |
|  | 1:0.75 |  |  | 0 | 0.01 |  |
|  | 0.75:1 |  |  | 0.01000 | 0.00625 |  |
|  | 1:0.5 |  |  | 0 | 0 |  |
| 3 | 1:1 | 78.98 | -0.0058 | 0 | 0.00400 | 0.47 |
|  | 1:0.75 |  |  | 0 | 0.01100 |  |
|  | 0.75:1 |  |  | 0.01 | 0.01000 |  |
|  | 1:0.5 |  |  | 0 | 0.01350 |  |
| 4 | 1:1 | 86.20 | -0.0485 | 0.00410 | 0.00740 | 0.51 |
|  | 1:0.75 |  |  | 0.00600 | 0.00510 |  |
|  | 0.75:1 |  |  | 0.01300 | 0 |  |
|  | 1:0.5 |  |  | 0.00150 | 0.00550 |  |

Figure 9 presents the fiber orientation distributions of the direct fiber model and the model’s mechanical anisotropy at maximum strain state. The findings suggest that the stiffness along each direction is approximately proportional to the amount of fibers in the loading direction at the maximum strain state.

**Figure 9.**
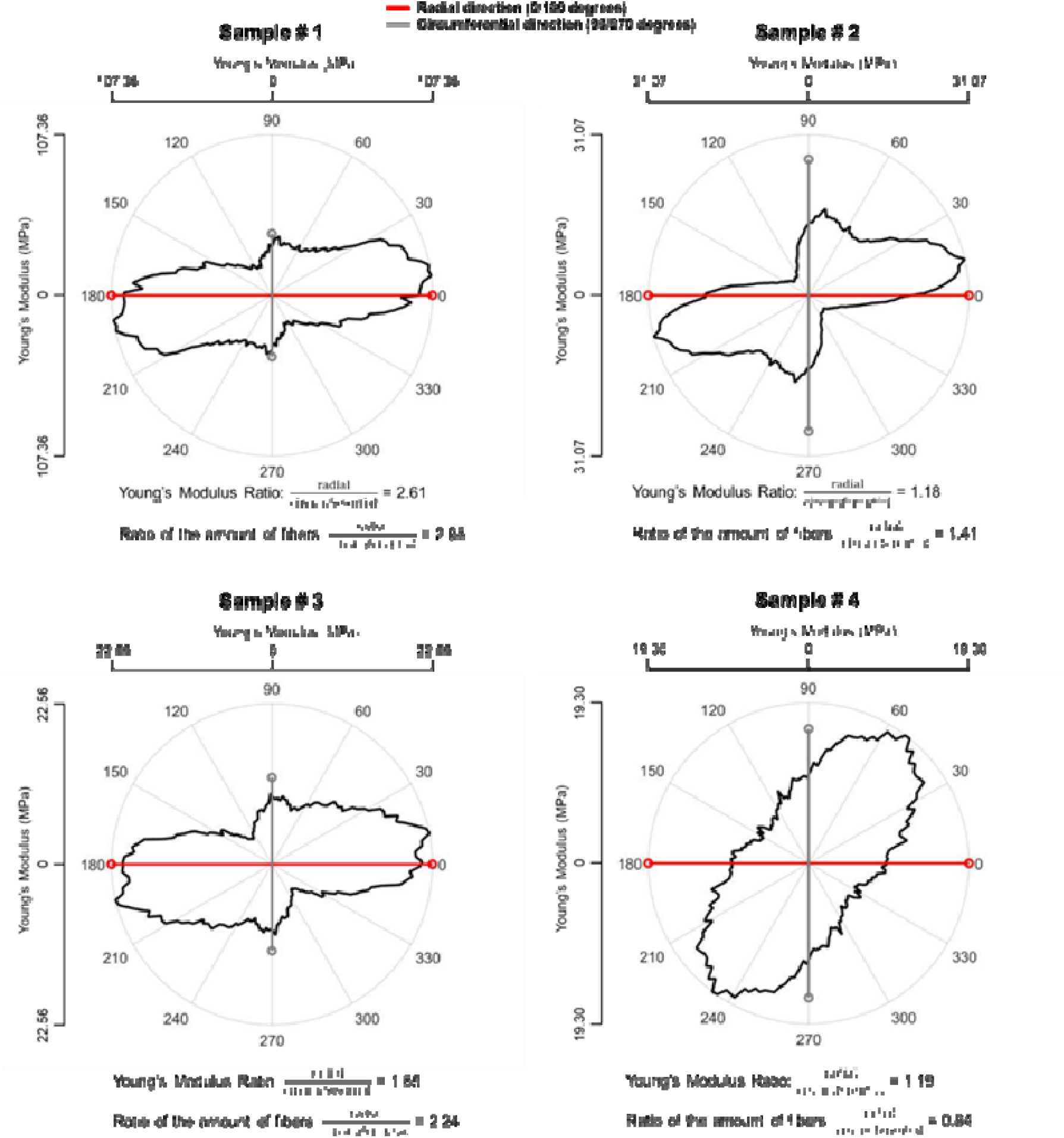
Orientation distribution and mechanical anisotropy of all the models. The orientation distributions are depicted in polar plots. Young’s modulus of each sample was estimated by calculating the slope of the model’s stress-strain curve at maximum strain state. The results indicate that at maximum strain state, the stiffness along each direction is approximately, but not exactly proportional to the amount of fibers aligned in the loading direction.

## 4. Discussion

Our goal was to continue advancing direct fiber models of soft tissues. Specifically, we aimed to conduct a more challenging test of the ability of the direct fiber modeling approach to capture sclera microstructure and anisotropic mechanics. We developed direct fiber models to simulate four sclera samples, incorporating specimen-specific three-dimensional fiber orientation distributions. Subsequently, we conducted an inverse fitting study and matched the models with specimen-specific anisotropic stress-strain behaviors from biaxial testing. The study yielded two main findings. First, the direct fiber models successfully captured the collagen structure of multiple sclera samples from different quadrants and eyes. Second, the macroscopic mechanical properties of the models matched with the experimental stress-strain data obtained under various anisotropic loading conditions. Notably, this was achieved by having fit the models to the equi-biaxial experiment. The derived material properties were then appropriate to represent the other loading conditions. This indicates that the direct fiber models inherently incorporated the anisotropy of tissue mechanical behaviors within their fiber structure, thereby eliminating the necessity for separate optimization with different loading conditions. Below we discuss the findings in detail.

### Finding 1. The direct fiber models can accurately capture the collagen fiber structure across different samples

In our previous study, the primary focus was on introducing the direct fiber modeling methodology, which involved utilizing experimental data from different samples and species obtained for other research purposes [17]. In this study, we deliberately chose to construct models that simulate multiple sclera samples from different quadrants and eyes. This decision was driven by the fact that the collagen fiber architecture of the sclera varies both spatially between locations and between different eyes [29]. Furthermore, it is known that the mechanical behavior of the sclera exhibit varying degrees of anisotropy across different quadrants, such as superior and temporal quadrants [41, 42]. By employing samples from both superior and temporal quadrants of multiple eyes, our study successfully demonstrated the robustness of the direct fiber model reconstruction approach in capturing the varying fiber structure. It is important to emphasize that our models were specimen-specific, meaning that the fiber structure was individually constructed based on each sample, and the model’s behaviors were optimized using experimental data obtained from the same sample.

### Finding 2: The macroscopic mechanical properties of the models matched with the experimental stress-strain data obtained under multiple anisotropic loading conditions

In our previous study, we constructed a single model and validated it against a specific equi- biaxial experimental dataset. In this study, we set the much tougher task of matching the stress-strain behaviors under multiple loading conditions. Our study thus indicates that the direct fiber modeling technique can account not only for equi-biaxial, but for the more complex anisotropic conditions that the sclera is subjected to [2]. We would like to point out here that we were unable to find examples of studies deriving material properties of sclera that simultaneously match multiple loading conditions. To the best of our knowledge, the standard seems to be to fit single experimental tests [43, 44]. Capturing tissue behavior under multiple loading conditions is a tougher challenge.

### Finding 3. The direct fiber models inherently incorporate the anisotropy of tissue mechanical behavior within their fiber structure

We have shown that the material properties obtained by fitting the equi-biaxial loading conditions were also adequate to produce good predictions (adjusted R^2^ > 0.9) for other biaxial loading tests. This indicates that the model reconstruction must have captured not only the tissue microarchitecture, as noted above in Finding 1, but also the anisotropic mechanical behavior encoded in the microarchitecture. We argue that these two, structural and mechanical anisotropies, while related, are not identical. Collagen fibers are the primary load-bearing component of the sclera, and therefore capturing sclera mechanical anisotropy requires a good representation of the microarchitecture [11, 44]. However, this may not be sufficient. The mechanical behavior, and particularly the nonlinear components, are heavily dependent on fiber undulations at multiple scales, and fiber-fiber interactions.[45, 46] Thus, we have shown a strength of the direct fiber modeling approach compared with the conventional continuum mechanical approach that requires inverse fitting several loading conditions [24]. While we acknowledge that our method is still an approximation of the actual fibrous structure of the sclera (more on this later in the subsection on Limitations), we argue that it represents a step forward for specimen-specific modeling.

### Interpretation of the derived fiber material properties

In our study, we initially derived the fiber material properties by matching the experimental data of loading protocol 1:1. Concerns may arise regarding the extent to which the results would be different if the material properties were obtained by matching experimental data from other loading protocols. To address this concern, we randomly selected a sample and derived the fiber material properties by independently fitting the loading protocols 1:0.75, 0.75:1, and 1:0.5. The fiber material properties were initially assigned random values and iteratively optimized to match the experimental data. Importantly, the fitting processes were performed without any reference to the results obtained from other loading protocols. Interestingly, we observed that the derived fiber material properties obtained by fitting different loading protocols fell within a similar range, as shown in Figure 10. To further evaluate the physical significance of these properties, we examined the resulting fiber elastic modulus and bulk modulus. Remarkably, these two properties demonstrated consistency across the results obtained from fitting different loading protocols. These findings emphasize the robustness of our method and confirm that the derived material properties are independent of the choice of loading protocols for fitting.

**Figure 10.**
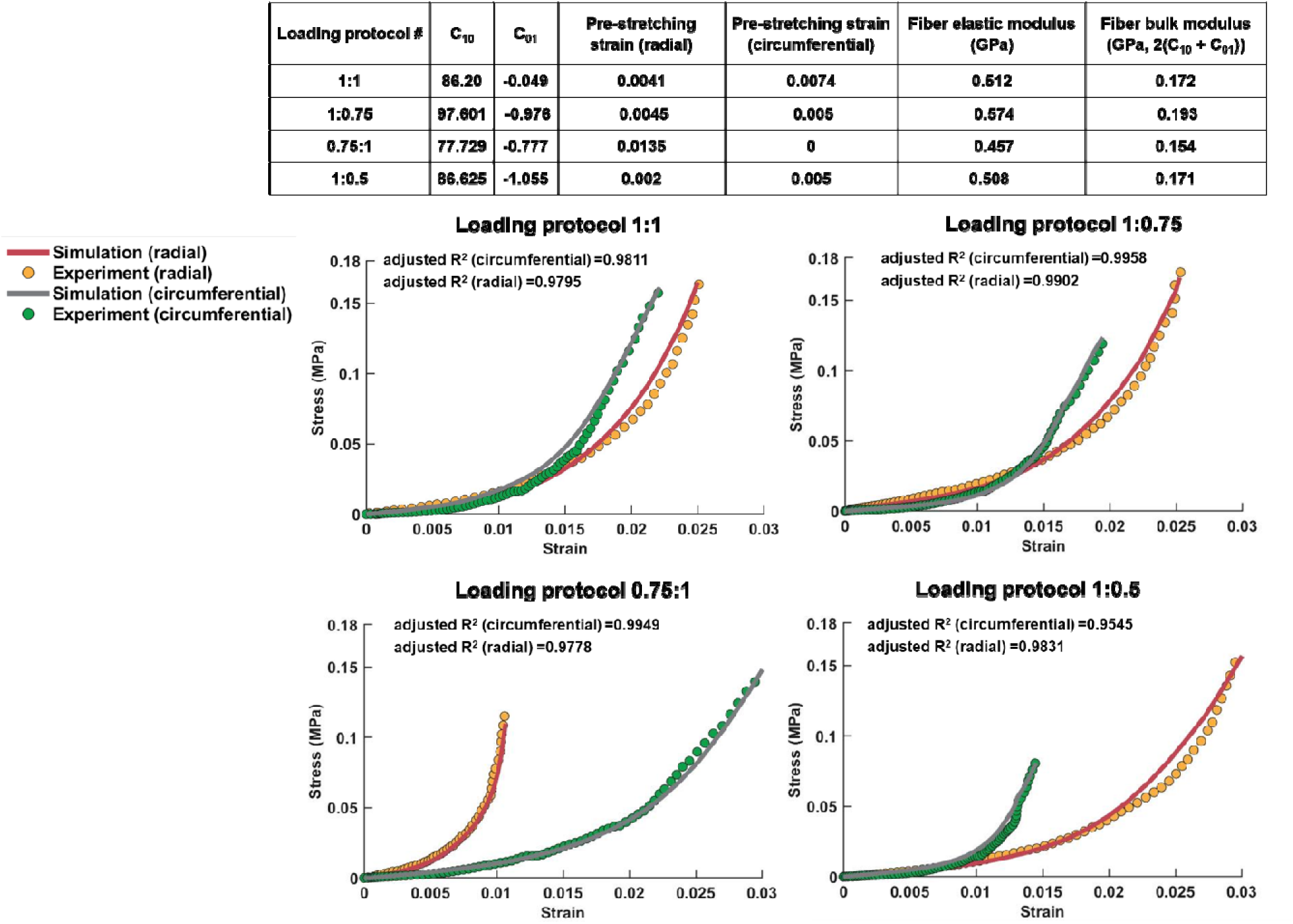
This figure illustrates the findings from the analysis using Sample #4 to assess the similarity of derived material properties when fitting the model to experimental data from different loading protocols. The table demonstrates that the values of C_10_ and C_01_ derived from fitting the model to different loading protocols are closely aligned, resulting in comparable fiber elastic modulus and fiber bulk modulus. Additionally, the stress-strain curves between the model and experimental data exhibit a strong overall fit, with an adjusted R^2^ greater than 0.9. These results support the conclusion that the choice of loading protocol for fitting does not influence the derived fiber material properties.

The “uniqueness” of so-called optimal parametes is a common concern in inverse fitting techniques [10, 11]. To address this concern, we conducted the inverse fitting process using four different sets of starting parameters to assess the consistency and uniqueness of the results. The derived fiber material properties, C_10_ and C_01_, exhibited significant variation among the four samples. To better understand those values which lack specific physical interpretation, we further estimated the elastic modulus of individual fibers by simulating uniaxial stretch of a single straight fiber with the optimal C_10_ and C_01_ values of the hyperelastic Mooney-Rivlin material. The estimated fiber elastic modulus ranged from 0.47 GPa to 23.94 GPa, and the detailed values can be found in Table 3. This wide range of values aligned with the experimentally reported elastic modulus values found in the literature, which ranged from 0.2 GPa to 7 GPa in bovine Achilles tendon fibers [47, 48], and 5 GPa to 11.5 GPa in rat tail [49]. It is worth noting that one value fell outside this range, which we believe may be attributed to two factors. Firstly, the load was primarily borne by a small portion of the fibers, leading to an overestimation of the fiber material properties. Secondly, the volume fraction of the models in this study was approximately 15%, which may not accurately represent the actual tissue composition. The estimated fiber material properties were closely associated with the fiber volume fraction in the model reconstructed. The difficulty of reconstructing models increases with the fiber density, as it becomes harder to follow the fibers in the histolory, and then reconstruct them accurately with their turns and undulations. Further research is necessary to comprehensively understand how variations in fiber volume fraction influence the behavior and derived material properties of the direct fiber model.

Another concern pertaining to the derived material properties and pre-stretching strains is the possibility of achieving an excessively favorable match between the models and experimental data across all loading protocols. To address this concern, we conducted a specific analysis using the fiber structure of Sample #1 and attempted to match the experimental data of sample #2. The results revealed that we could successfully match the experimental data for loading protocols 1:1, 1:0.75, and 0.75:1. However, when attempting to match the experimental data for loading protocol 1:0.5, the model could not achieve a satisfactory fit (Figure 11). This indicates that the model is not universally applicable but rather depends on accurately matching the orientation distribution of the sample in order to replicate the experimental data of that specific sample.

**Figure 11.**
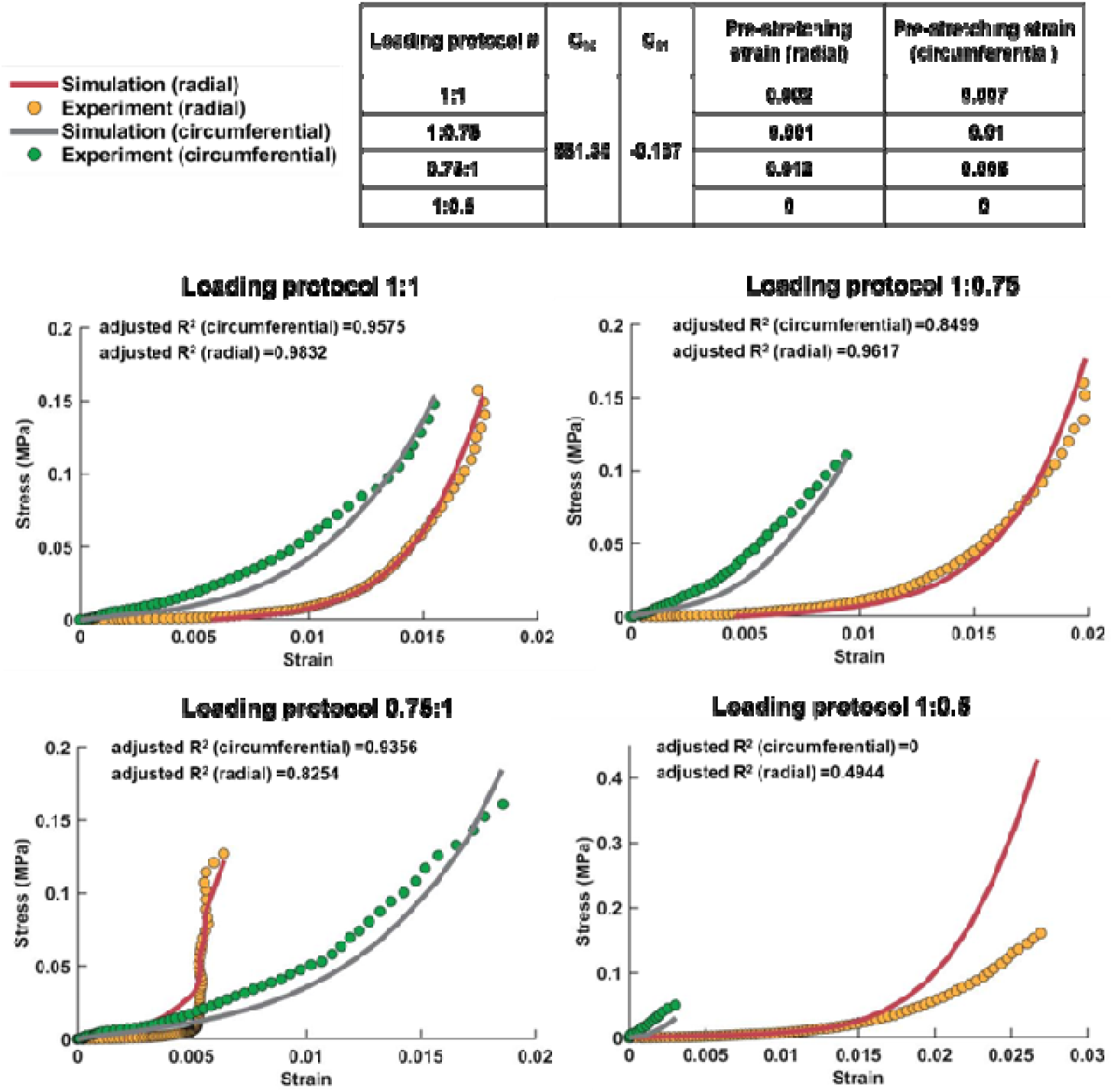
This figure presents the results of using the model fiber structure of Sample #1 to fit the experimental stress-strain data of Sample #2, aimed at assessing the possibility of achieving an overly favorable match between the models and experimental data. Following the process described in the main text, the fiber material properties C10 and C01 were derived by fitting the model to loading protocol 1:1. The results indicate successful matches between the model and experimental data for loading protocols 1:1, 1:0.75, and 0.75:1, with overall adjusted R^2^ values exceeding 0.8. However, for loading protocol 1:0.5, the model exhibits softer behavior than the experiment in the circumferential direction and stiffer behavior in the radial direction, resulting in adjusted R^2^ values lower than 0.5. As such, it cannot be considered as a valid match. These findings reinforce that the model’s applicability is not universal but rather dependent on accurately matching the orientation distribution of the specific sample in order to replicate its experimental data.

### Strengths of direct fiber models

The models developed in this study inherited several strengths, some of which we have discussed above or elsewhere [17]. Briefly, the direct fiber models have considered fiber interweaving and the resulting fiber-fiber interactions that play an important role in determining the structural stiffness of the sclera [15]. The models also have included collagen fibers of the sclera that are long and continuous. Thus, they can transfer forces over a long distance [2, 14], where it’s commonly recognized that it’s important to account for long fiber mechanics [19, 50–52]. This is in contrast to the conventional continuum mechanics approaches where fibers exist only within a given element, transferring their loads at the element boundaries [53]. Furthermore, the direct fiber models in this work incorporate three-dimensional specimen-specific collagen architecture, whereas previous fiber-aware models tend to overlook or simplify the in-depth orientations of the collagen fibers [42, 54, 55]. Collagen fiber variations in-depth of the tissue are crucial in determining the sclera’s load-bearing capacity, particularly in bearing shear stresses, and may have clinical implications [56, 57].

In comparison to our preliminary study, we implemented two improvements to the approach when building direct fiber model structures. Firstly, we incorporated the natural curvature of the tissue into the model, enabling a more accurate representation of the physiological shape of the sclera. This enhancement ensured that the model mimicked more closely the three-dimensional geometry of the sclera, improving its fidelity. Secondly, we introduced a constraint function in Abaqus to deal with fibers that were not sufficiently constrained. This helped avoid, for example, cases where a fiber could be “pulled out” of the model. This constraint function served to tightly constrain the interconnections between fibers. As a result, the model not only exhibited increased stability, but also better matched the actual interweaving and cohesion of collagen fibers in the sclera. This optimization helped overcome the challenge of inaccuracies caused by floating fibers and enhances the model’s reliability and validity in representing the real tissue structure. We suspect that this constraint would not be necessary if we had modeled in more detail the fiber-fiber and fiber-matrix interactions. The simplifications we assumed on these interactions allow fibers much freedom to displace and slide. It is also possible that our modeling of somewhat thick fiber bundles instead of small scale fibers may not have captured the full extent of the fiber entanglement that prevents fiber sliding. Further work is necessary to characterize these aspects of the sclera and how to account for them accurately in models. Herein we just want to comment that we acknowledge that our treating of fiber-fiber and fiber-matrix interactions, while limited, is still more comprehensive than that in conventional continuum models where these are generally not only ignored but cannot be incorporated without cumbersome kludges. Our direct fiber modeling approach highlights the assumptions on the interactions and provides a platform for improving their modeling.

## Limitations

The first limitation we would like to comment on was observed in the matching of stress-strain curves between the model and experimental data. While the majority of loading conditions exhibited strong agreement with adjusted R^2^ values exceeding 0.9, there were instances, particularly in the loading protocol 1:0.5, where the model’s fit was relatively low. We do not know the cause, but this discrepancy may result from the sequential application of five different loading protocols during the testing process. After multiple rounds of testing, the tissue may become softer than its original condition. As a result, the model’s behavior appeared to be stiffer than the corresponding experimental data in these cases. Future studies should either randomize the loading protocols, or explicitly study the effects of protocol order. Despite this limitation, it is important to emphasize that the direct fiber modeling technique demonstrated robustness and effectiveness in capturing the overall mechanical behaviors of the sclera, as evidenced by the strong agreement between the model and experimental data in the majority of the cases. The observed discrepancies in certain loading conditions provide valuable insights for future refinements of the model and highlight the need for further investigation into the dynamic changes in tissue properties during sequential loading.

Second, we encountered difficulties in accurately matching the loading protocol 0.5:1 due to the model’s inability to perform well under compression. This limitation indicated that further optimization of the approach is required to enable stable simulations under compression. Future efforts should be focused on addressing this issue to enhance the applicability of the direct fiber modeling approach. By improving the model’s performance in compression scenarios, we can broaden its utility in studying the mechanical behaviors of the sclera across a wider range of loading conditions.

Third, our study used sheep eyes as the sample model, and thus the question arises as to how well this would work with other tissues, or with other species. Considering the robustness of the approach in capturing the complex fiber structure and anisotropic mechanical behaviors of sheep sclera, we anticipated that the direct fiber modeling technique can serve as a valid and effective tool for investigating scleral biomechanics in other species and potentially in other collagen-based tissues as well. Future studies can extend the application of this approach to validate its broader utility and ensure its broad application across various tissues.

Fourth, the direct fiber models, while consistent with experimentally measured fiber orientation distribution, are still approximations of the actual tissue. Several aspects were simplified or ignored in the modeling process, which may introduce discrepancies between the models and the real tissue. One such simplification was the assumption of uniform fiber or bundle diameters within the models. In reality, collagen fibers in the sclera can exhibit variations in diameter [58–60]. Additionally, sub-fiber level features were not explicitly included in the models, such as fiber crimp [5]. Future work would benefit from incorporating more detailed and realistic microstructural features, where the models can provide a more accurate representation of the scleral tissue and enhance our understanding of its mechanical behavior.

Fifth, the matrix mechanical properties were kept constant at literature values and not optimized iteratively like the fiber properties. This simplification was made for simplicity. Although the matrix properties could potentially affect fiber load-bearing and parameter fitting, their impact is considered minor. The fibers, being the primary load-bearing component, exhibit significantly greater stiffness compared to the matrix [11, 44]. Analysis of our model indicated that the matrix bears only 4%–6% of the total reaction forces at the maximum strain. Therefore, we believe that the fiber properties predominantly influence the model’s behavior, while the matrix stiffness has minimal influence on the derived parameters. In future work, it would be worthwhile to explore the role of matrix properties and consider more detailed matrix-fiber interactions for further refinement.

Sixth, we ignored fiber-matrix interactions and considered simplified fiber-fiber interactions, similar to our previous work [17]. However, it should be noted that the interactions between fibers and the matrix can be much more complex and may influence the tissue behaviors [61, 62]. Future research could benefit from incorporating more realistic and complex interactions within the direct fiber modeling framework. The direct fiber modeling technique has the potential to be a valuable tool for studying and considering such complex interactions in future studies.

In conclusion, we performed a comprehensive and challenging test of the direct fiber modeling approach through the simulation of multiple sheep posterior sclera samples. We characterized the macroscopic and anisotropic stress-strain behaviors of the samples through biaxial mechanical testing. Then direct fiber models were generated based on the microstructural architecture of each sample. An inverse fitting process was employed to simulate biaxial stretching conditions, enabling the determination of optimal pre-stretching strains and fiber material properties. Our findings demonstrated the efficacy of the direct fiber modeling approach in simulating the scleral microarchitecture, capturing critical fiber characteristics, and accurately describing its anisotropic macroscale mechanical behaviors. Moreover, we highlighted the capability of the direct fiber models to inherently incorporate the anisotropy of tissue mechanical behaviors within their fiber structure. Overall, the direct fiber modeling approach proved to be a robust and effective tool for characterizing the biomechanics of sclera.

## Supplemental data

**Figure 1.**
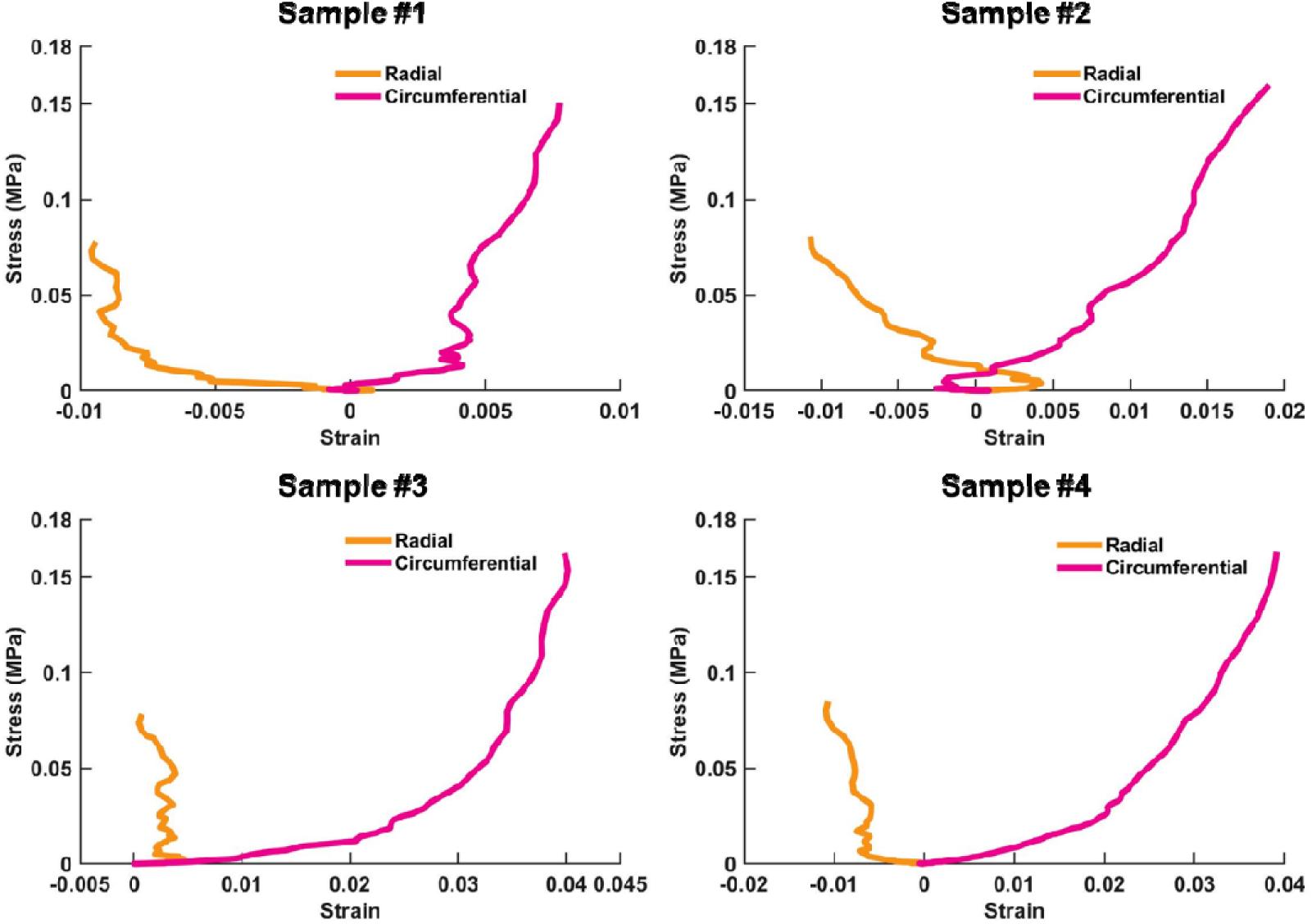
Stress-strain curves of the loading protocol 0.5:1 of the four samples. Under this loading condition, where the circumferential direction experienced higher stress compared to the radial direction, the radial direction of the sample exhibited a contraction, leading to negative strains.

